# LRBA promotes drug-induced liver injury and MASLD by scaffolding MAPK activation

**DOI:** 10.1101/2025.11.17.688803

**Authors:** Daisuke Oikawa, Min Gi, Yoshinori Okina, Duc Pham Minh, Hidetaka Kosako, Le Thi Thanh Thuy, Yusuke Sato, Hiroko Ikenaga, Mana Kosugi, Tsutomu Matsubara, Kouhei Shimizu, Masayuki Shiota, Fumiaki Ando, Shinichi Uchida, Hirotaka Takahashi, Takuro Horii, Izuho Hatada, Kazuo Ikeda, Tatsuya Sawasaki, Norifumi Kawada, Fuminori Tokunaga

## Abstract

LPS-responsive *beige*-like anchor protein (LRBA) regulates vesicular trafficking and receptor recycling, and its deficiency results in immunodeficiency characterized by hypogammaglobulinemia and autoimmune syndrome. However, its role in liver pathophysiology remains unclear. Here, we reveal a previously unrecognized function of LRBA as a critical intracellular scaffold for mitogen-activated protein kinase (MAPK) activation that promotes liver injury. *Lrba^-/-^*mice exhibit reduced acetaminophen (APAP)-induced hepatic necrosis through the suppression of JNK activation. In a model of metabolic dysfunction-associated steatotic liver disease (MASLD) induced by a high-fat, high-cholesterol (HFHC) diet, *Lrba* deficiency reduces hepatic inflammation, fibrosis, and Kupffer cell activation. Mechanistically, LRBA homodimers directly interact with specific mitogen-activated protein kinase kinase kinases (MAP3Ks), including transforming growth factor-β-activated kinase 1 (TAK1) and mixed-lineage kinase 3 (MLK3), to facilitate their activation. LRBA, which is primarily expressed in hepatocytes under physiological conditions, is upregulated in non-parenchymal cells such as Kupffer cells and cholangiocytes in both HFHC diet-fed mice and patients with MASLD and cirrhosis, linking its scaffolding function to pathological inflammation. Thus, LRBA promotes liver disease progression by amplifying inflammatory signaling.

## Introduction

Lipopolysaccharide (LPS)-responsive *beige*-like anchor protein (LRBA) is encoded by an LPS-inducible gene and exhibits characteristics of A-kinase anchoring proteins (AKAPs) and *beige* Chediak-Higashi syndrome (BEACH) domain-containing proteins (BDCPs) (Wang *et al*, 2001). *LRBA* encodes a ∼319 kDa (2,863 amino acids) endosomal protein involved in vesicular trafficking and receptor recycling (Martinez Jaramillo & Trujillo-Vargas, 2018). Mutations or deficiencies in *LRBA* are associated with common variable immunodeficiency (CVID) (Lopez-Herrera *et al*, 2012), a disorder characterized by recurrent infections beginning in childhood, hypogammaglobulinemia, and various autoimmune manifestations, including idiopathic thrombocytopenic purpura, autoimmune hemolytic anemia, inflammatory bowel disease (IBD), and chronic erosive polyarthritis (OMIM #606453) (Levy *et al*, 2016).

Although *Lrba^-/-^* mice do not exhibit the hypogammaglobulinemia characteristic of CVID, they show increased susceptibility to dextran sulfate sodium (DSS)-induced colitis (Wang *et al*, 2019). Interestingly, *LRBA* deficiency can be ameliorated by abatacept, a cytotoxic T lymphocyte antigen-4 (CTLA-4)-immunoglobulin fusion protein, as LRBA regulates the intracellular recycling of CTLA-4 from endosomes to the plasma membrane (Lo *et al*, 2015) *via* interactions with Rab11 (Janman *et al*, 2021). Additionally, the Arf1-dependent recruitment of LRBA to Rab4-positive endosomes is critical for maintaining endolysosomal homeostasis (Szentgyorgyi *et al*, 2024). We have also shown that LRBA functions as an AKAP to facilitate the phosphorylation of aquaporin 2 (AQP2), thereby contributing to urinary concentration in the kidney (Hara *et al*, 2022; Yanagawa *et al*, 2023).

Souza *et al*. employed computational approaches to predict that *LRBA* is one of several previously uncharacterized genes that exacerbate drug-induced liver injury (DILI) (Souza *et al*, 2018). They also proposed that these so-called “dark genes” could be involved in liver diseases beyond DILI. Since the role of LRBA in liver pathophysiology remains unclear, we investigated its function in various disease models using *Lrba*^-/-^ mice. Our findings demonstrate that *Lrba* deficiency ameliorates liver injury across multiple models, including acetaminophen (APAP)-induced DILI and high-fat, high-cholesterol (HFHC) diet-induced metabolic dysfunction-associated steatotic liver disease (MASLD). Mechanistically, LRBA forms homodimers and acts as a scaffold for specific mitogen-activated protein kinase kinase kinases (MAP3Ks), such as transforming growth factor-β-activated kinase 1 (TAK1, also known as MAP3K7) and mixed-lineage kinase 3 (MLK3, also known as MAP3K11), thereby facilitating their activation. Consequently, *Lrba* deficiency suppressed the activation of stress-activated protein kinases (SAPKs), including c-Jun N-terminal kinase (JNK). Collectively, these findings suggest that the LRBA-mediated regulation of inflammatory signaling represents a potential therapeutic target for the treatment of liver diseases, including MASLD.

## Results

### *Lrba* deficiency ameliorates DILI

Recent computational analyses suggested that LRBA may be a risk factor for DILI caused by APAP, diclofenac, isoniazid, nimesulide, and valproic acid (Souza *et al*., 2018). However, the association between LRBA and liver disease remains unclear. To investigate the role of LRBA in DILI, we administered APAP to wild-type (*Lrba*^⁺/⁺^) and *Lrba*^-/-^ mice and evaluated the resulting acute hepatotoxicity. Histological analyses with hematoxylin and eosin (H&E) staining and necrosis scoring revealed a marked reduction in APAP-induced centrilobular hepatocellular necrosis in *Lrba*^-/-^ mice (Fig. 1A, B). This was accompanied by significantly lower serum levels of alanine aminotransferase (ALT) and aspartate aminotransferase (AST), as well as attenuated expression of inflammatory genes, compared with *Lrba*^⁺/⁺^ mice (Fig. 1C, D).

**Figure 1.**
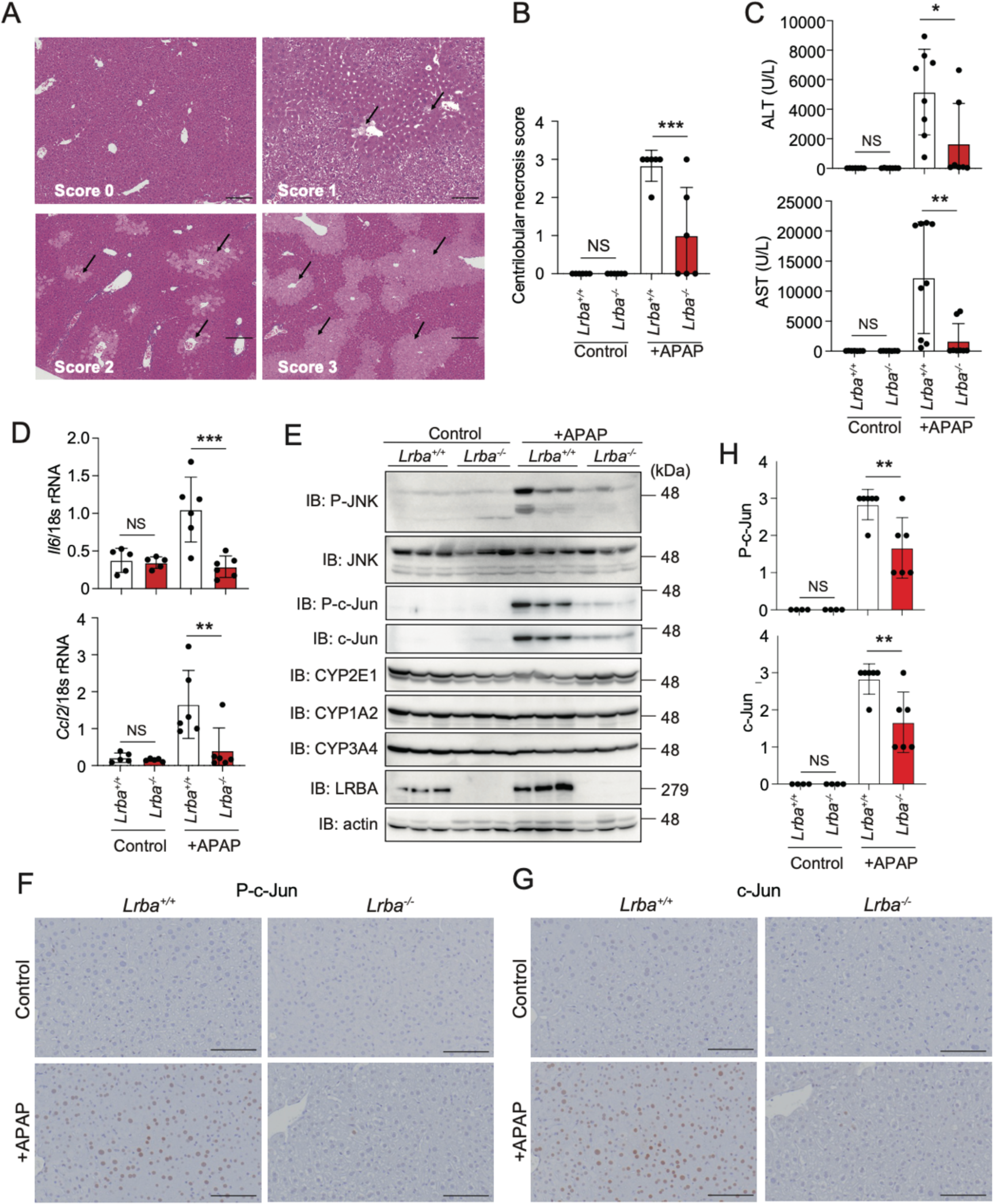
Suppressed APAP-induced liver injury in *Lrba*^-/-^ mice. (**A**) Representative images of centrilobular hepatocellular necrosis at each score level, 6 h after intraperitoneal (i.p.) injection of 300 mg/kg APAP. Score 0 (normal): *Lrba*^+/+^ mouse injected with saline (control); score 1 (minimal): *Lrba***^-/-^** mouse treated with APAP; score 2 (moderate): *Lrba***^-/-^** mouse treated with APAP; score 3 (severe): *Lrba*^+/+^ mouse treated with APAP. *Bars*, 200 μm. (**B**) Significantly lower scores of centrilobular hepatocellular necrosis in APAP-treated *Lrba***^-/-^**mice compared to APAP-treated *Lrba*^+/+^ mice (*n* = 6). (**C**) Reduced ALT and AST levels in APAP-injected *Lrba***^-/-^** mice. Serum ALT and AST levels were measured in *Lrba*^+/+^ and *Lrba***^-/-^** mice (*n* = 7-9) with or without APAP injection. (**D**) Suppressed induction of inflammatory genes in APAP-treated *Lrba^-/-^* mice. qPCR analysis was performed using total liver RNA collected 6 h after APAP injection (*n* = 5-6). (**E**) Decreased activation of JNK and c-Jun in livers from APAP-injected *Lrba***^-/-^**mice. Immunoblotting was performed on liver lysates from *Lrba*^+/+^ and *Lrba***^-/-^** mice injected with or without APAP (3 per condition), using the indicated antibodies. (**F**-**H**) Reduced activation of c-Jun in APAP-treated *Lrba***^-/-^** mice. Representative liver images showing P-c-Jun (**F**) or c-Jun staining (**G**), and their quantification (**H**) (*n* = 4-6). *Bars*, 100 µm. Data are presented as individual values with mean ± SEM. *p < 0.05, **p < 0.01, ***p < 0.001 by the ANOVA post-hoc Tukey test (B-D, and H). NS, not significant.

During DILI, APAP is metabolized by cytochrome P450 enzymes, such as CYP2E1, into the metabolite *N*-acetyl-*p*-benzoquinone imine (NAPQI), which generates reactive oxygen species (ROS) that activate JNK, thereby promoting hepatotoxicity, cell death, and inflammation (Nakagawa *et al*, 2008; Seki *et al*, 2012). In *Lrba*^-/-^ mice, the APAP-induced activation of JNK and its downstream target c-Jun was reduced, despite the unchanged expression levels of CYP enzymes (Fig. 1E-H). These findings suggest that LRBA plays a critical role in APAP-induced DILI by regulating activation of the mitogen-activated protein kinase (MAPK) signaling pathway.

### LRBA promotes HFHC diet-fed MASLD phenotypes

To further investigate the role of LRBA in liver pathophysiology, we employed an HFHC diet-fed MASLD model. After 40 weeks of feeding with either a normal chow (NC) or an HFHC diet, hepatomegaly, liver weight, and serum levels of ALT and AST were reduced in *Lrba*⁻/⁻ mice compared with *Lrba*⁺/⁺ mice, whereas their body weights were comparable (Figs. 2A and EV1A-C). H&E staining and MASLD activity scoring indicated attenuated liver damage in *Lrba*^-/-^ mice (Fig. 2A, B). Immunohistochemical analyses revealed reduced fibrosis, decreased activation of fibrosis-associated factors, and diminished T cell infiltration in HFHC-fed *Lrba*^-/-^ mice (Fig. 2A, B). qPCR analyses further confirmed the markedly suppressed expression of genes associated with fibrosis and inflammation in *Lrba*^-/-^ mice (Fig. 2C, D).

**Figure 2.**
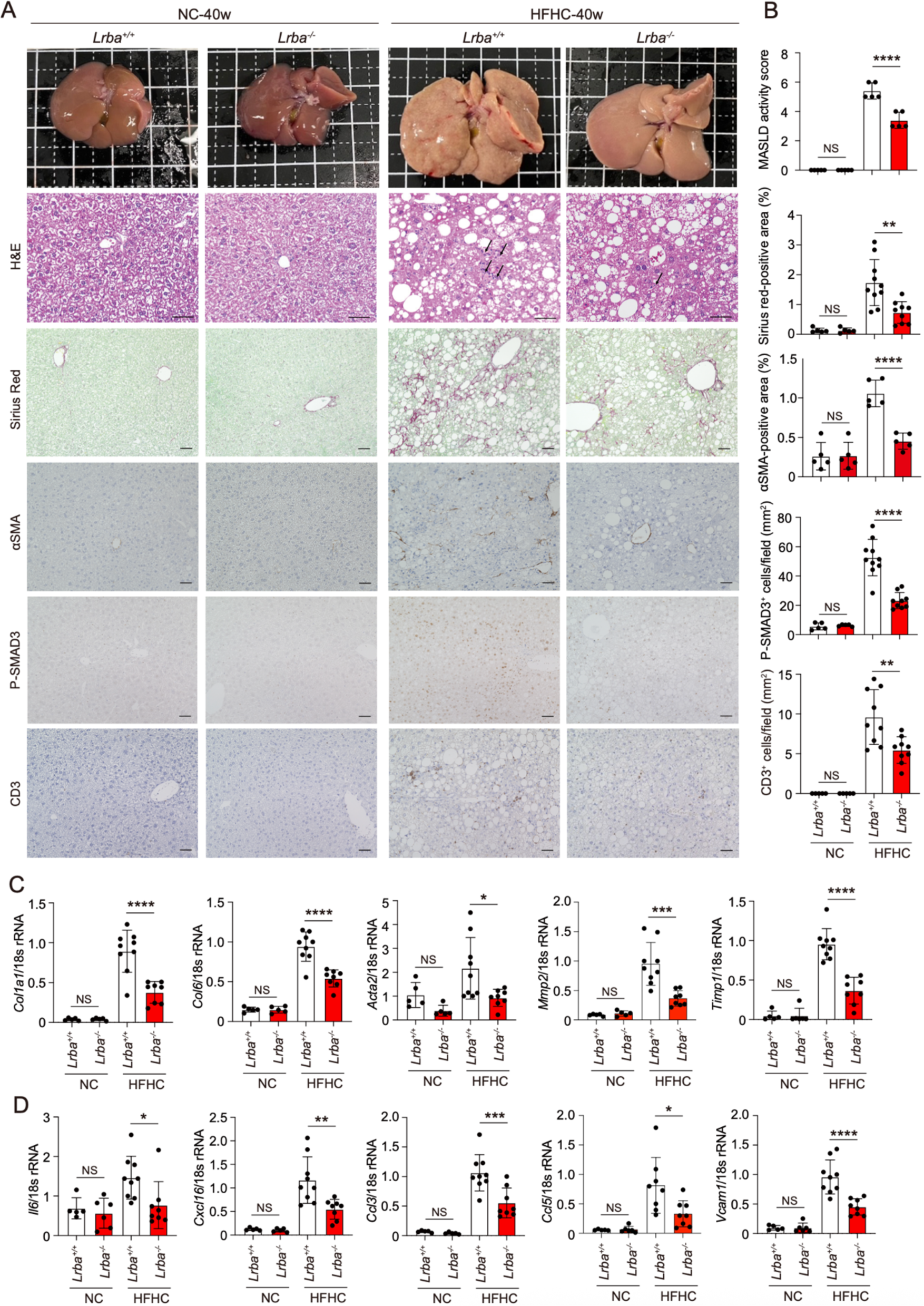
HFHC diet-induced MASLD is attenuated in *Lrba*^-/-^ mice. (**A**, **B**) Attenuated hepatic fibrosis and inflammation in HFHC-fed *Lrba*^-/-^ mice. *Lrba*^+/+^ and *Lrba*^-/-^ mice were fed either NC- or HFHC-diet for 40 weeks. Representative macroscopic liver images, H&E staining, Sirius Red staining, and immunohistochemistry for αSMA, P-SMAD3, and CD3 are shown (**A**), along with quantification of staining intensities and positive areas (**B**) (*n* = 5-10). *Bars*, 50 µm. *Arrows* in H&E: infiltrated immune cells. (**C**, **D**) Reduced expression of fibrogenic and inflammatory genes in HFHC-fed *Lrba*^-/-^ mice. qPCR analysis of fibrosis-related genes (**C**) and inflammatory-related genes (**D**) was performed using total liver RNA (*n* = 5-9). Data are presented as individual values with mean ± SEM. *p < 0.05, **p < 0.01, ***p < 0.001, ****p < 0.0001 by the ANOVA post-hoc Tukey test (B-D). NS, not significant.

In contrast, no significant differences were observed in Oil Red O-positive lipid accumulation or lipid concentrations in the serum and liver between HFHC-fed *Lrba*^⁺/⁺^ and *Lrba*^-/-^ mice (Fig. EV1D-F). Although lipid peroxidation, as indicated by 4-hydroxynonenal (4-HNE) staining, was reduced in HFHC-fed *Lrba*^-/-^ mice (Fig. EV1G, H), the expression of oxidative stress-related genes remained unchanged (Fig. EV1I). These results indicate that *Lrba* deficiency suppresses HFHC diet-induced MASLD primarily by attenuating inflammatory and fibrotic pathways, rather than by altering lipid accumulation.

### Kupffer cell activation is attenuated in HFHC-fed *Lrba*^-/-^ mice

To investigate the mechanisms underlying the attenuated MASLD phenotype in *Lrba*^-/-^ mice, we performed single-cell fixed RNA sequencing (scRNA-seq) using frozen liver sections. Uniform Manifold Approximation and Projection (UMAP) analysis identified nine distinct liver cell clusters (Fig. EV2A-C). In hepatocyte clusters, enrichment analysis revealed that acute inflammatory responses and positive regulation of the ERK1/ERK2 cascade were attenuated in HFHC-fed *Lrba*^-/-^ mice compared with *Lrba*^+/+^ mice (Fig. EV2D, E). In hepatic stellate cell (HSC) clusters, collagen and fibril organization was diminished in *Lrba*^-/-^ mice (Fig. EV2G, H). In macrophages, signatures of myeloid cell activation, protein autophosphorylation, and positive regulation of the JNK cascade were attenuated (Fig. EV2J, K). In contrast, gene sets related to xenobiotic metabolic processes, steroid metabolism, and oxidative phosphorylation were upregulated in HFHC-fed *Lrba*^-/-^ mice (Fig. EV2F, I, L). Intercellular signaling analyses further suggested that the TGF-β and EphA signaling pathways were suppressed in HFHC-fed *Lrba*^-/-^ mice, whereas protease-activated receptor (PAR) signaling was upregulated (Fig. EV2M, N).

Importantly, scRNA-seq analysis revealed that the macrophage cluster was further subdivided into four distinct subclusters, among which activated Kupffer cells (aKUPs) were markedly expanded in HFHC-fed *Lrba*⁺/⁺ mice but substantially reduced in *Lrba*⁻/⁻ mice (Fig. 3A, B). The expression of aKUP marker genes, including *Trem2* and *Gpnmb*, was significantly decreased in HFHC-fed *Lrba*^-/-^ mice compared with *Lrba*^+/+^ mice (Fig. 3C). Notably, cell-cell interactions involving macrophage-derived Trem2-Tyrobp signaling were lost in *Lrba*^-/-^ mice (Fig. 3D). Consistent with these findings, immunoblotting and qPCR analyses using total liver lysates and RNAs, respectively, demonstrated the reduced expression of components associated with the Trem2-Tyrobp pathway in *Lrba*^-/-^ mice (Fig. 3E, F). Immunohistochemical analysis further confirmed that GPNMB, a well-established marker of aKUPs (McGettigan *et al*, 2019; Xiong *et al*, 2019), was sparsely expressed in macrophages distributed throughout the hepatic sinusoids in control *Lrba^⁺/⁺^* and *Lrba^⁻/⁻^* mice, without hepatocellular fat degeneration or clustering (Fig. 3 G). By contrast, in HFHC diet-fed *Lrba^⁺/⁺^* and *Lrba^⁻/⁻^* mice, GPNMB-positive macrophage clusters were localized around steatotic hepatocytes. Notably, the abundance of these clusters was markedly reduced in *Lrba^⁻/⁻^* mice (Fig. 3G). The GPNMB-positive macrophage cluster scores were significantly reduced in HFHC diet-fed *Lrba^⁻/⁻^* mice compared with HFHC diet-fed *Lrba^⁺/⁺^* mice (Fig. 3H). Immunofluorescence analysis further confirmed that GPNMB expression was enriched in CLEC4F-positive Kupffer cells in *Lrba*^+/+^ mice, whereas the portion of GPNMB-positive Kupffer cells was significantly lower in *Lrba^⁻/⁻^* mice (Fig. 3I, J). Collectively, these results indicate that *Lrba* deficiency suppresses the inflammatory activation of Kupffer cells *via* the Trem2-Tyrobp pathway, in the context of HFHC-induced liver injury.

**Figure 3.**
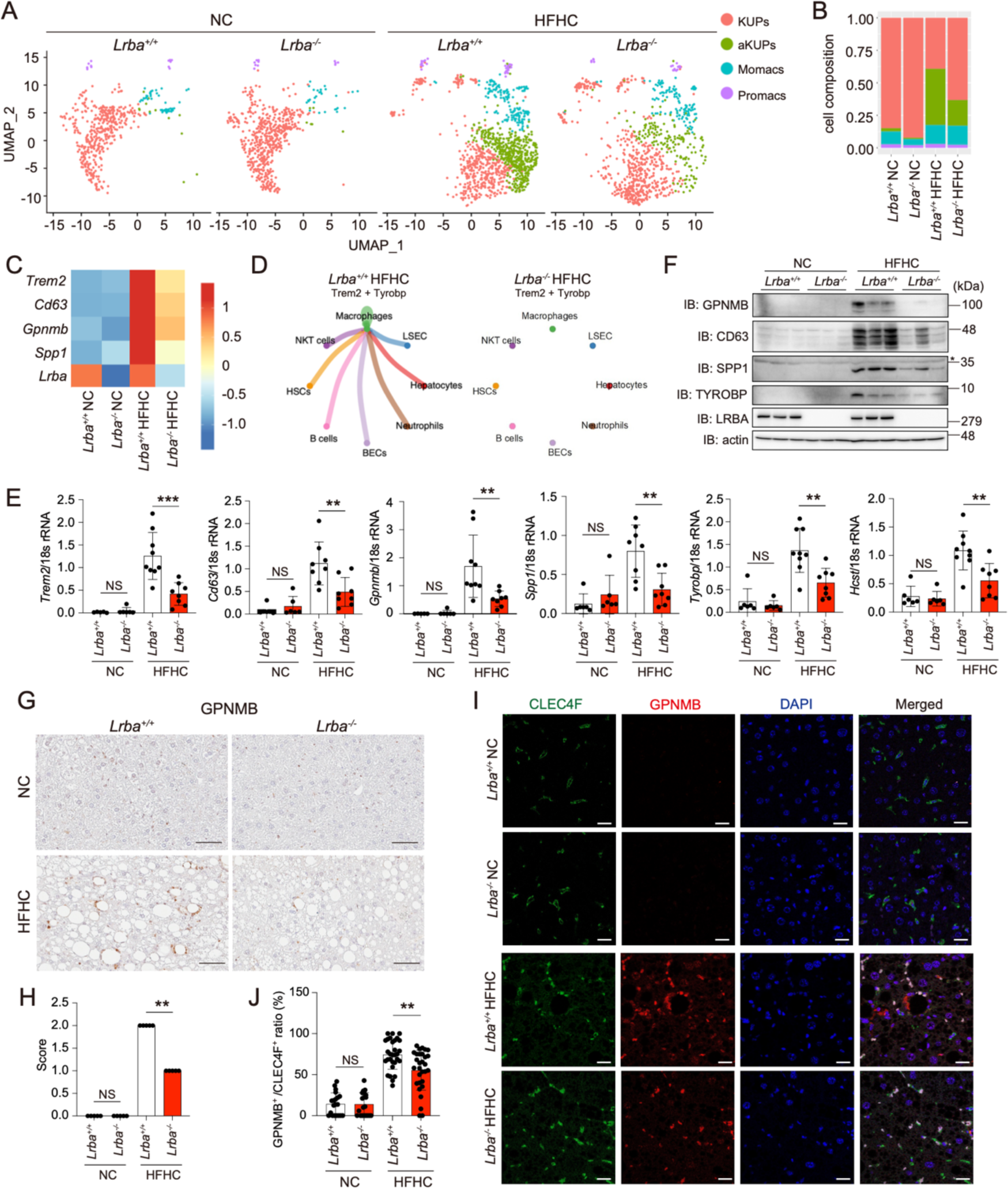
Kupffer cell activation is attenuated in HFHC-fed *Lrba*^-/-^ mice. (**A**-**D**) scRNA-seq analysis of liver macrophage populations. UMAP plots with four distinct macrophage subclusters (**A**), the proportional composition of each subcluster (**B**), a heatmap showing marker gene expression in aKUPs (**C**), and the cell-cell interactions mediated by the Trem2-Tyrobp signaling pathway (**D**) are shown. (**E**, **F**) Decreased expression of aKUP markers in HFHC-fed *Lrba*^-/-^ mice. qPCR analysis of Trem2-Tyrobp pathway components using total liver RNA (**E**) (*n* = 5-9) and immunoblotting of liver lysates from triplicate samples using the indicated antibodies (**F**) were performed. *: non-specific signal. (**G**, **H**) Reduced accumulation of GPNMB-positive cells in livers of HFHC-fed *Lrba*^-/-^ mice. Representative images of GPNMB staining (**G**) and its quantification (**H**) are indicated. *Bars*, 60 µm. (**I**, **J**) Kupffer cell activation was attenuated in *Lrba*^-/-^ mice. Immunofluorescent staining of the Kupffer cell markers CLEC4F (green) and GPNMB (red), nuclei (DAPI), and their merged images (**I**), and quantification of the ratio of GPNMB^+^ / CLEC4F^+^ cells (**J**) (*n* = 20-34). *Bars*, 20 μm. Data are presented as individual values with mean ± SEM. **p < 0.01, ***p < 0.001 by the ANOVA post-hoc Tukey test (E, H, J). NS, not significant.

### MAPK signaling activation is attenuated in *Lrba*-deficient cells

Given that TLR4-mediated MAPK signaling, which also crosstalks with the Trem2-Tyrobp pathway, is a key driver of MASLD-associated Kupffer cell activation (Kazankov *et al*, 2019; Miura & Ohnishi, 2014), we examined the role of LRBA in this pathway using the mouse Kupffer cell line KUP5 (Kitani *et al*, 2014). Knockdown of *Lrba* did not alter the protein or mRNA levels of Trem2-Tyrobp pathway components or TLR4 (Fig. EV3A, B). However, *Lrba* knockdown attenuated LPS-induced activation of the MAPK and NF-κB signaling pathways, as well as the expression of their downstream target genes, in KUP5 cells (Fig. 4A, B). Similarly, bone marrow-derived macrophages (BMDMs) from *Lrba*^-/-^ mice showed reduced MAPK activation (Fig. 4C). Moreover, stimulation of BMDMs with various TLR ligands or agonists resulted in suppressed induction of target genes in *Lrba*^-/-^ cells (Fig. 4D). Likewise, LPS- or IL-1β-induced activation of MAPK was diminished in *Lrba*^-/-^ mouse embryonic fibroblasts (MEFs), and restoration of LRBA in *Lrba*^-/-^ MEFs rescued MAPK activation (Fig. EV3C, D). Furthermore, lysosomal stress-induced activation of JNK, which has been implicated in Kupffer cell activation (Kanamori *et al*, 2021), was also attenuated in *Lrba*^-/-^ MEFs (Fig. EV3E). Collectively, these results indicate that LRBA is essential for MAPK signaling activation, and that its loss compromises both innate immune sensing and inflammatory responses.

**Figure 4.**
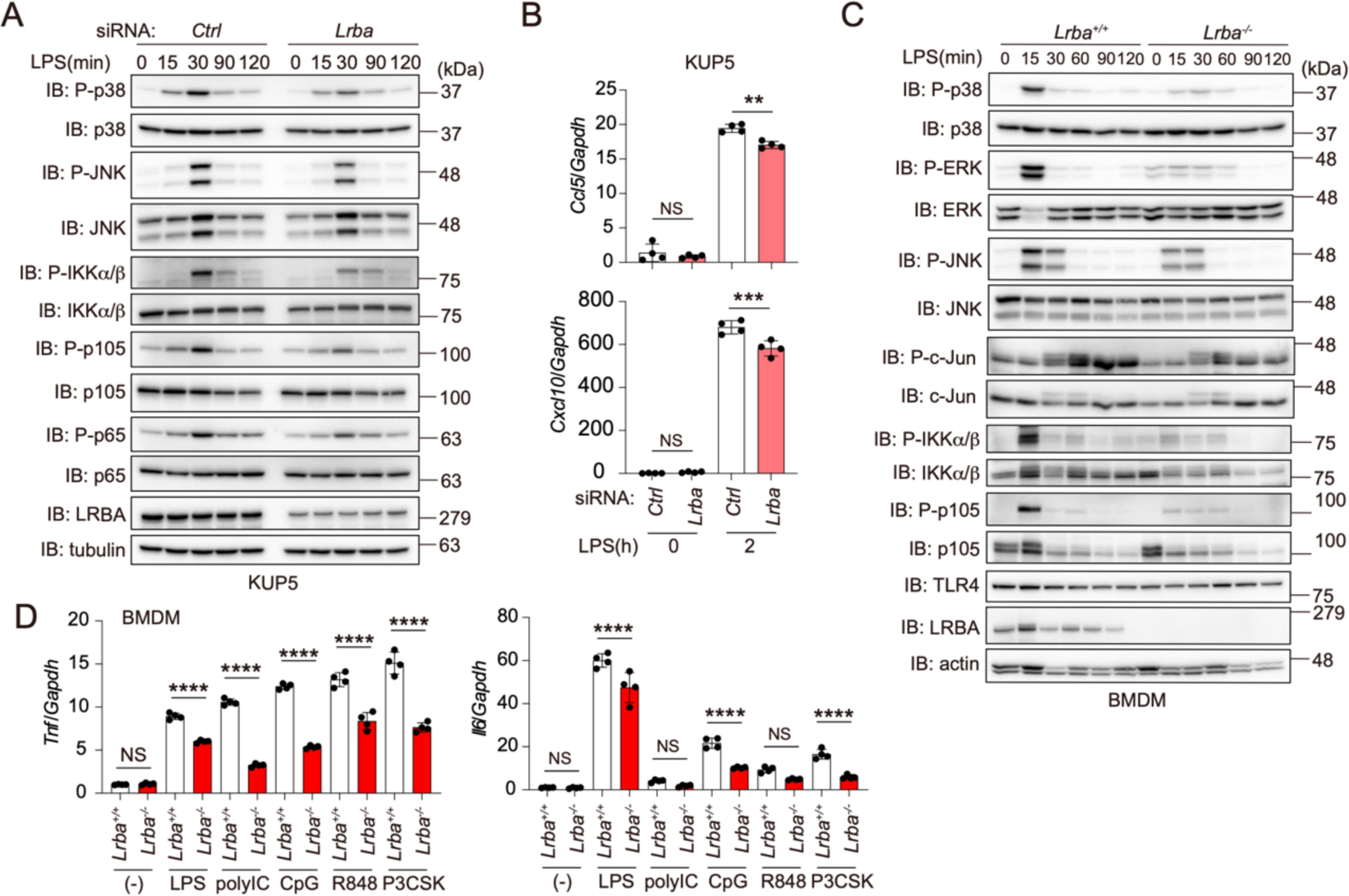
Suppressed activation of MAPK signaling in *Lrba*-deficient cells. (**A**, **B**) Knockdown of *Lrba* attenuates MAPK signaling in a Kupffer cell line. KUP5 mouse Kupffer cells were transfected with the indicated siRNAs and treated with 100 ng/ml LPS for the indicated periods. Cell lysates were subjected to immunoblotting with the indicated antibodies (**A**) or qPCR analysis (**B**) (*n* = 4). (**C**) LPS-stimulated MAPK activation is attenuated in *Lrba***^-/-^** BMDMs. BMDMs from *Lrba*^+/+^ and *Lrba***^-/-^** mice were stimulated with 100 ng/ml LPS for the indicated periods and analyzed by immunoblotting. (**D**) Attenuated expression of PAMPs-induced inflammatory genes in *Lrba***^-/-^** BMDMs. BMDMs were stimulated with LPS or R848 for 4 h, or with polyI:C, CpG, or Pam3CSK4 for 8 h, and qPCR analysis was performed (*n* = 4). Data are presented as individual values with mean ± SEM. **p < 0.01, ***p < 0.001, ****p < 0.0001 by the ANOVA post-hoc Tukey test (B, D). NS, not significant.

### LRBA dimer functions as a scaffold for TAK1 and MLK3 to activate MAPK signaling

To elucidate the molecular basis by which LRBA regulates MAPK signaling, we performed proximity labeling using AirID-tagged LRBA followed by mass spectrometry (Fig. EV4A; Appendix Table S1). A volcano plot revealed self-labeling of LRBA, suggesting intermolecular association (Fig. EV4B). Gel filtration chromatography of lysates from *Lrba*^+/+^ and *Lrba*^-/-^ MEFs revealed that endogenous LRBA, despite a predicted monomeric mass of ∼309 kDa, eluted in a fraction corresponding to ∼669 kDa, suggesting dimer formation in cells (Fig. EV4C). Co-immunoprecipitation analysis provided further confirmation, as Myc-tagged LRBA co-precipitated with FLAG-LRBA (Fig. EV4D).

Among the MAPK-related proteins identified by proximity labeling, we identified interactions with MEK1, MEKK1, and TAK1-binding protein 2 (TAB2), a scaffold protein required for TAK1 activation (Takaesu *et al*, 2000) (Fig. EV4B). However, co-immunoprecipitation analysis showed negligible interactions between LRBA and either MEK1 or MEKK1 (Fig. EV4E). In contrast, LRBA bound directly to TAK1 and associated with the TAK1-TAB2 complex (Fig. 5A), and this interaction was enhanced upon IL-1β or LPS stimulation (Fig. 5B). Truncation mapping identified the LRBA PH-BEACH domain (Gebauer *et al*, 2004) (a.a. 2045-2483) as responsible for TAK1 binding (Fig. 5C, D). AlphaFold 3 modeling predicted that this domain interacts with the N-terminal kinase domain of TAK1 through a specific interface involving LRBA Phe2406 and TAK1 Phe237 (Figs. 5E and EV4F). Consistent with this model, the interaction was abolished in the TAK1 F237A and LRBA F2406R mutants (Fig. 5F, G). Further AlphaFold 3-based screening predicted MLK3 (MAP3K11) and MAP3K19 as additional candidate interactors among MAPK/IKK family kinases (Fig. EV4G, H). Co-immunoprecipitation confirmed that LRBA associates with MLK3, but not with the kinase domain of MAP3K19 (a.a. 1055-1328) (Fig. 5H). The endogenous LRBA-MLK3 interaction was enhanced following LPS stimulation (Fig. 5I). Supporting a model in which TAK1 and MLK3 bind the PH-BEACH domain in distinct orientations, wild type (Wt)-LRBA interacted with Wt-MLK3, but not with MLK3 mutants (E121A, V122R, or E121A/ V122R) or the LRBA F2465R mutant (Fig. EV4I, J).

**Figure 5.**
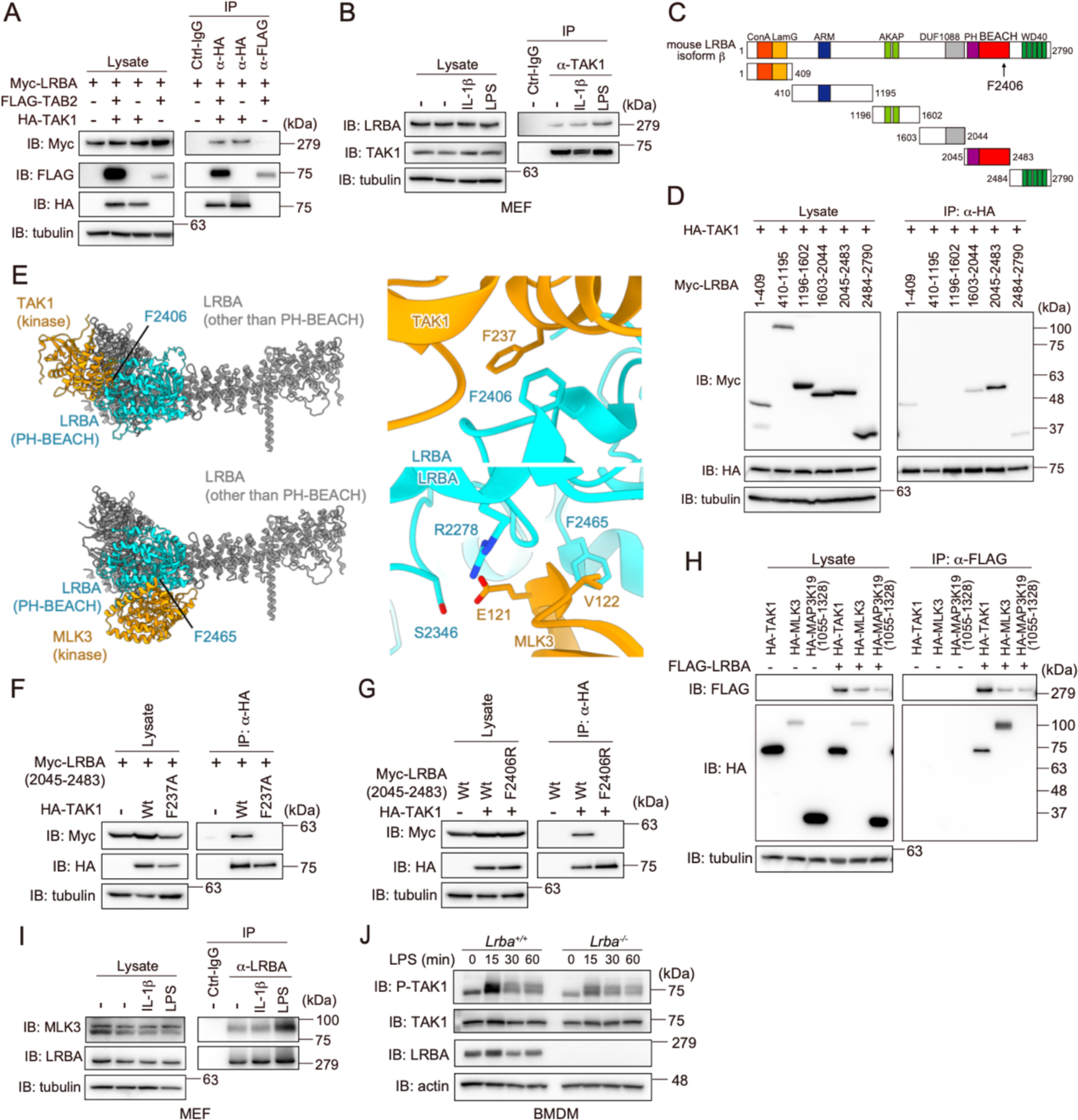
LRBA interacts with TAK1-TAB2/3 and MLK3 to activate MAPK pathway. (**A**) LRBA interacts with TAK1. FLAG-TAB2 and HA-TAK1 were co-expressed with Myc-LRBA in HEK293T cells. Cell lysates and anti-HA or anti-FLAG-immunoprecipitates were immunoblotted with the indicated antibodies. (**B**) Endogenous association between LRBA and TAK1 is enhanced upon stimulation. *Lrba*^+/+^ MEFs were stimulated with 20 ng/ml IL-1β for 10 min or 5 μg/ml LPS for 20 min. Cell lysates and anti-TAK1 immunoprecipitates were analyzed by immunoblotting. (**C**) Schematic representation of mouse LRBA domain structure and truncated fragments. ConA: ConA-like/lectin domain; LamG: laminin G-like domain; ARM: armadillo/β-catenin-like repeats; AKAP: PKA/C-binding motifs; DUF: a conserved domain of unknown function; PH: Pleckstrin homology domain; BEACH: *beige* and Chediak-Higashi syndrome domain. (**D**) The PH-BEACH domain (a.a. 2045-2483) of LRBA interacts with TAK1. Myc-tagged LRBA fragments were co-expressed with HA-TAK1 in HEK293T cells. Cell lysates and anti-HA-immunoprecipitates were immunoblotted with the indicated antibodies. (**E**) Structural models of LRBA PH-BEACH domain in complex with TAK1 (*top*) or MLK3 (*bottom*) predicted by AlphaFold 3. PH-BEACH domain: *cyan*; other LRBA regions: *grey*; kinase domains of TAK1 and MLK3: *orange*. *Left*, full-length LRBA superimposed with TAK1 or MLK3; *right*, close-up views of the BEACH–kinase interaction interfaces. Flexible regions are omitted for clarity. (**F**) Phe237 in TAK1 is critical for LRBA-binding. Co-immunoprecipitation was performed with Wt- or F237A mutant of HA-TAK1 and Myc-LRBA (a.a. 2045-2483). (**G**) Phe2406 in LRBA is critical for TAK1-binding. Co-immunoprecipitation analysis was performed using Wt- or F2406R mutant of Myc-LRBA (a.a. 2045-2483) and HA-TAK1-Wt. (**H**) LRBA interacts with MLK3. A similar co-immunoprecipitation analysis as in (A) was performed using FLAG-LRBA co-expressed with HA-TAK1, HA-MLK3, or HA-MAP3K19 (a.a. 1055-1328) in HEK293T cells. (**I**) Endogenous interaction between LRBA and MLK3. Co-immunoprecipitation was performed using the anti-LRBA antibody as in (B). (**J**) TAK1 activation is impaired in *Lrba*^-/-^ BMDMs. BMDMs from *Lrba*^+/+^ and *Lrba*^-/-^ mice were stimulated with 100 ng/ml LPS, and lysates were subjected to immunoblotting.

Consistently, LPS-induced TAK1 activation was impaired in *Lrba*^-/-^ BMDMs (Fig. 5J), although the association between TAK1 and TAB2/TAB3 remained unchanged in *Lrba*^+/+^ and *Lrba*^-/-^ MEFs (Fig. EV4K). Notably, TAB3 immunoprecipitated with TAK1 exhibited an electrophoretic mobility shift following IL-1β or LPS treatment, likely reflecting phosphorylation (Mendoza *et al*, 2008), and this shift was reduced in *Lrba*^-/-^ MEFs (Fig. EV4K). Together, these findings indicate that LRBA dimerizes in cells and acts as a scaffold for TAK1 and MLK3, thereby facilitating MAPK pathway activation.

### Increased LRBA expression in cholangiocytes is associated with MASLD

LRBA is broadly expressed across multiple organs and tissues (Roussa *et al*, 2024). In the livers of NC diet-fed mice, LRBA was predominantly detected in albumin-positive hepatocytes (Fig. 6A). In contrast, LRBA expression was markedly upregulated in non-parenchymal cells that partially overlapped with CLEC4F-positive Kupffer cells and cytokeratin 19 (CK19)-positive cholangiocytes in HFHC diet-fed mice (Fig. 6B). This upregulation was corroborated by a total liver RNA analysis, which showed elevated *Lrba* transcript levels following HFHC feeding (Fig. 6C). These results suggest that MASLD progression in mice is associated with increased LRBA expression in non-parenchymal cell populations, particularly Kupffer cells and cholangiocytes.

**Figure 6.**
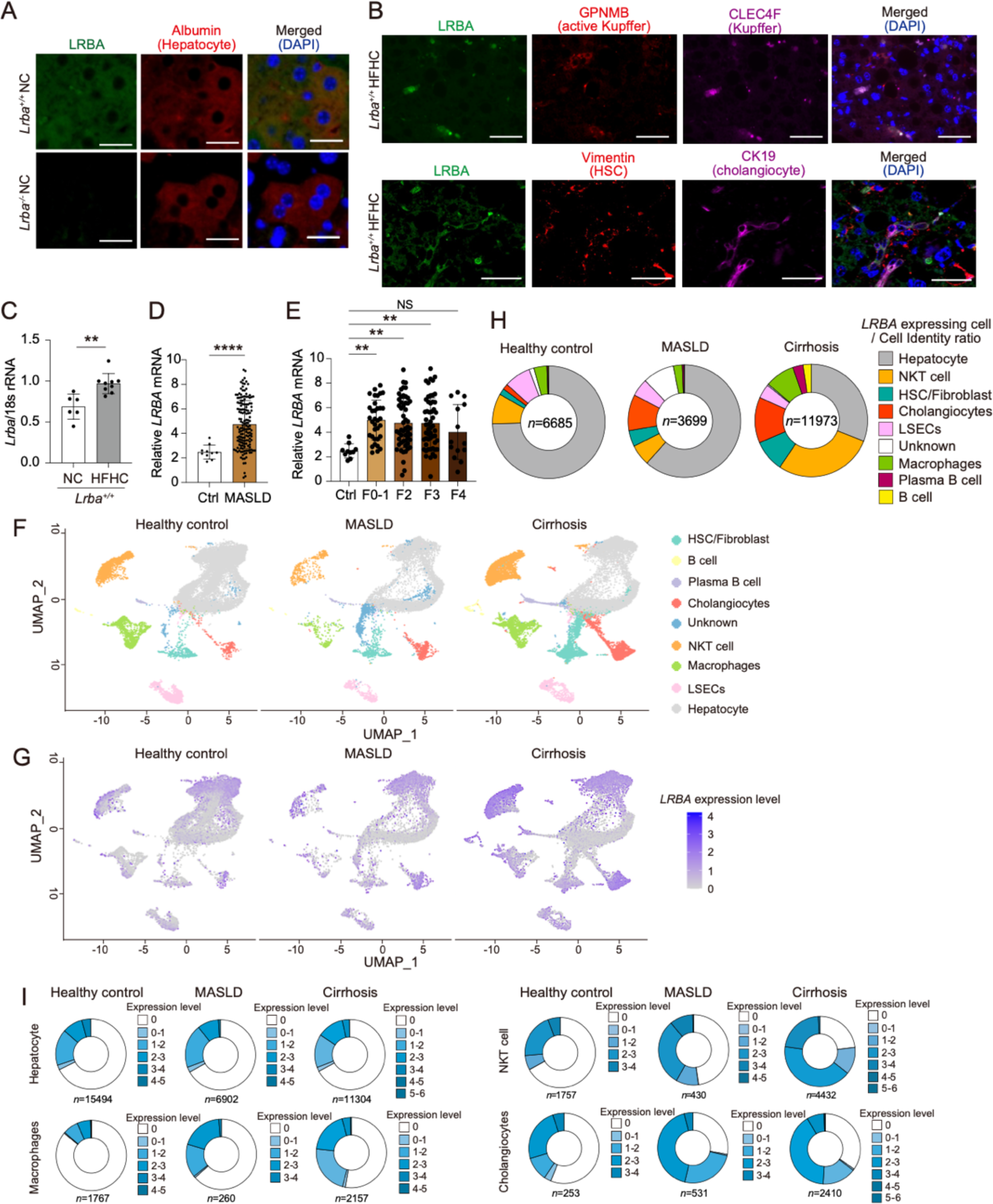
*LRBA* is upregulated in non-parenchymal cells, such as cholangiocytes, in patients with MASLD and cirrhosis. (**A**) LRBA expression in hepatocytes under the NC conditions. Representative immunofluorescence images of liver sections from NC-fed *Lrba*^+/+^ and *Lrba*^-/-^ mice stained for LRBA (*green*), cell-type-specific markers (*red*), and merged images are shown. *Bars*, 20 μm. (**B**) LRBA expression is upregulated in Kupffer cells and cholangiocytes of HFHC diet-fed *Lrba*^+/+^ mice. A similar analysis as in (A) was performed using liver sections from HFHC-fed *Lrba*^+/+^ mice. *Bars*, 20 μm. (**C**) Induction of *Lrba* mRNA upon HFHC diet-feeding. qPCR analysis was performed on total liver RNA from NC- and HFHC-fed mice (*n* = 6, 9). (**D**, **E**) *LRBA* mRNA expression is elevated in liver tissues from MASLD patients. RNAseq data analysis of liver tissues was used to compare *LRBA* expression between healthy controls and MASLD patients (**D**) and across MASLD disease stages (**E**). (**F-I**) LRBA is predominantly expressed in non-parenchymal liver cells from patients with MASLD and cirrhosis. UMAP plot displaying the nine distinct liver cell clusters from scRNA-seq data of healthy controls, MASLD, and cirrhosis patients (**F**), UMAP plot highlighting *LRBA*-expressing cells (**G**), proportions of each cluster expressing *LRBA* (**H**), and LRBA expression levels across each cluster (**I**) are shown. Data are presented as individual values with mean ± SEM. \*\**p* < 0.01, **** *p* < 0.0001 by the Mann-Whitney test (C, D) or the ANOVA post-hoc Tukey test (E). NS, not significant.

To assess *LRBA* expression in human liver disease, we analyzed publicly available bulk transcriptome datasets. *LRBA* expression was upregulated in MASLD patients from the early fibrotic stages, although its levels did not progressively increase with fibrosis severity (Fig. 6D, E). Further analysis using public scRNA-seq data from healthy controls and individuals with MASLD or cirrhosis revealed that *LRBA* was primarily expressed in hepatocytes under healthy conditions (Fig. 6F-H). In contrast, *LRBA* expression was markedly increased in non-parenchymal cells in both MASLD and cirrhotic livers (Fig. 6F-H). Notably, *LRBA* levels in hepatocytes remained unchanged between healthy and diseased states, whereas *LRBA* expression in non-parenchymal cells was significantly elevated in both MASLD and cirrhosis (Fig. 6I). These findings are consistent with observations from the murine MASLD model and indicate that LRBA expression in non-parenchymal cells plays a key role in the pathogenesis of human MASLD and liver cirrhosis.

## Discussion

LRBA, a ubiquitously expressed member of the BDCP family, has been implicated in a variety of diseases, including immunodeficiency (Burns *et al*, 2012; Levy *et al*., 2016; Lopez-Herrera *et al*., 2012), IBD (Alangari *et al*, 2012; Gettler *et al*, 2021; Wang *et al*., 2019), modulation of CTLA-4-targeted therapies (Lo *et al*., 2015), and the regulation of water reabsorption in the kidney through the control of AQP2 phosphorylation (Hara *et al*., 2022). Although LRBA had previously been predicted to be a risk factor for DILI (Souza *et al*., 2018), our study is the first to experimentally demonstrate that *Lrba* deficiency in mice protects against APAP-induced hepatotoxicity and mitigates MASLD progression under HFHC diet conditions.

We demonstrated that LRBA functions as a scaffold protein for MAP3Ks, such as TAK1-TAB2/3 and MLK3, in its homodimeric form, thereby facilitating the MAPK a signaling to promote inflammatory responses and cell death (Fig. EV5A). In APAP-induced DILI, apoptosis signal-regulating kinase 1 (ASK1), MKK4, and JNK1/2 act as MAP3K, MAP2K, and MAPK, respectively, and hepatotoxicity is linked to the prolonged activation of JNK1/2 (Nakagawa *et al*., 2008). Notably, this phenotype is attenuated in *Mlk3^-/-^* mice (Sharma *et al*, 2012). LRBA has also been proposed to act as an AKAP, sharing high homology with DAKAP550 and containing two putative PKA-binding motifs that interact with the regulatory subunit RIIβ of PKA (Moreno-Corona *et al*, 2020). Our structural and biochemical analyses showed that the TAK1- or MLK3-binding regions in the PH-BEACH domain of LRBA are distinct from the PKA-binding motif. Specifically, Phe2406 and Phe2465 within this domain are critical residues mediating TAK1 and MLK3 binding, respectively (Fig. 5). These residues are conserved across multiple BDCP family members (Fig. EV5B), suggesting a broad scaffolding function with this protein family. Furthermore, the evolutionary conservation of this regulatory mechanism is supported by the presence of LRBA orthologues in yeast (*Bph1*) and *Drosophila* (*rugose, rg*). Mutations in *rg* lead to a shortened lifespan and a rough-eye phenotype, mediated by JNK-dependent apoptosis (Wech & Nagel, 2005). Collectively, these findings suggest that the role of LRBA in MAPK signaling is evolutionarily conserved and represents a fundamental mechanism for the underlying stress and inflammatory responses.

The progression of MASLD is driven by a complex interplay of inflammatory responses, oxidative stress, endoplasmic reticulum (ER) stress, and insulin resistance. These pathogenic processes promote the transition from simple hepatic steatosis to fibrosis, cirrhosis, and ultimately hepatocellular carcinoma. Among the central regulators of these pathways are protein kinases, particularly those within the MAPK signaling cascade, which orchestrate key pathogenic events. Accordingly, the modulation of MAPK activity has emerged as a promising therapeutic strategy (Alshehade *et al*, 2022). ASK1, a member of the MAP3K family, is currently being explored as a drug target, with several selective inhibitors undergoing clinical investigation (Ray, 2017). In addition, hepatic phosphoglycerate dehydrogenase (PHGDH) has been shown to protect against MASLD by suppressing TAK1 activation (Hu *et al*, 2025). Interestingly, *Mlk3*^-/-^ mice exhibit substantial protection in MASLD models, which is characterized by reduced hepatic macrophage infiltration and activation (Gadang *et al*, 2013; Ibrahim *et al*, 2014). These phenotypes closely resemble those observed in *Lrba*^-/-^ mice, suggesting that the MLK3-LRBA axis may play a key role in the pathogenesis of MASLD.

The BDCP family comprises nine members: lysosomal trafficking regulator (LYST), neurobeachin (NBEA), neurobeachin-like 1 (NBEAL1), neurobeachin-like 2 (NBEAL2), LRBA, WD and FYVE zinc finger domain-containing proteins 3 and 4 (WDFY3 and WDFY4), neutral sphingomyelinase activation-associated factor (NSMAF), and WD repeat domain 81 (WDR81) (Cullinane *et al*, 2013). Among these, LRBA shares the highest amino acid sequence homology (∼75%) with NBEA. Genetic mutations in *NBEA* have been linked to neurodevelopmental disorders with early-onset generalized epilepsy (OMIM #604889) and are also associated with Alzheimer’s disease (Martinelli-Boneschi *et al*, 2013). Notably, the TLR4-dependent MAPK signaling pathway and TREM2 expression have been implicated in the pathogenesis of Alzheimer’s disease (Ruganzu *et al*, 2022), raising the possibility that NBEA, like LRBA, may function as a key regulator of neuroinflammation. Moreover, WDFY3 is highly expressed in the liver (Chen *et al*, 2004) and has been identified as a prognostic biomarker in long-term survivors of extrahepatic cholangiocarcinoma (Wang *et al*, 2020). These findings collectively suggest that BDCP family proteins may contribute to disease progression not only through their interactions with PKA, but also by functioning as scaffold proteins that modulate MAPK signaling. Thus, members of the BDCP family represent promising targets for future therapeutic development.

## Methods

### Mice

*Lrba*^-/-^ mice were generated in our laboratory, as previously described (Hara *et al*., 2022). All mice were cared for according to the guidelines approved by the Institutional Animal Care and Use Committee of Osaka Metropolitan University. In the DILI model, acetaminophen (*N*-acetyl-*p*-aminophenol: APAP) dissolved in isotonic saline at 65°C was administered intraperitoneally (i.p.) at a dose of 300 mg/kg to 8- to 12-week-old male mice, which were sacrificed 6 h later. For the MASLD model, 7-week-old male mice were fed either a high-fat, high-cholesterol (HFHC) diet or normal chow (NC) for 40 weeks. All diets were purchased from the Oriental Yeast Co., Ltd. (Tokyo, Japan), and their compositions are shown in Appendix Table S2. Serum and liver tissues were collected and stored at −80°C for subsequent analyses.

### ALT and AST assay

ALT and AST concentrations in mouse serum were quantified using the Transaminase CII test kit (FUJIFILM, Tokyo, Japan).

### Tchol, TG, and NEFA measurements

The liver tissues were homogenized in 50 mM Tris-HCl buffer, pH 7.5, containing 0.1% Triton X-100. The liver lysates and serum samples were analyzed for bile acid (BA), total cholesterol (TChol), triglyceride (TG), and nonesterified fatty acid (NEFA), using FUJIFILM Wako Pure Chemical assay kits (Wako, Japan).

### Histopathological analysis

Liver tissues were fixed in 4% paraformaldehyde (PFA), embedded in paraffin, and sectioned at 4 μm thickness. Sections were stained with hematoxylin and eosin (H&E), Sirius Red, or Oil Red O, or subjected to immunohistochemical analysis. Centrilobular hepatocellular necrosis induced by APAP was semi-quantitatively scored based on the extent of necrotic areas using the following criteria: 0, normal; 1, minimal; 2, moderate; 3, severe. The number of c-Jun- or phosphorylated c-Jun (P-c-Jun)-positive hepatocytes was evaluated using the following scoring system: 0: none; 1: <1% of hepatocytes; 2: positive cells ≤5% of hepatocytes; 3: ≥50% of hepatocytes. GPNMB-positive macrophage clusters were scored as follows: 0, none; 1, clusters surrounding <5% of hepatocytes with fatty degeneration; 2, clusters surrounding >5% of hepatocytes with fatty degeneration.

### Sirius red, Oil Red O, and immunohistochemical staining

For Sirius red staining, paraffin-embedded liver sections were stained with 0.1 % (w/v) Sirius Red (Direct Red 80) and 0.1% Fast Green FCF in a saturated aqueous picric acid solution. For Oil Red O staining, frozen liver sections were rinsed with distilled water, stained with 0.18% Oil Red O (Sigma-Aldrich) with 60% 2-propanol (Sigma-Aldrich) for 15 minutes at 37°C, and counterstained with hematoxylin. For immunohistochemical staining, paraffin sections were deparaffinized and dehydrated through a graded ethanol series, then heated for 20 min at 120°C in citrate acid buffer for antigen retrieval. Endogenous peroxidase activity was quenched using 0.3% hydrogen peroxide. After blocking with 3% bovine serum albumin (BSA) in phosphate-buffered saline (PBS) for 1 h, the sections were incubated with primary antibodies overnight at 4°C, and with secondary antibodies for 1 h at room temperature. Images were obtained using a BZ-X700 All-in-One fluorescence microscope (Keyence, Osaka, Japan), and positive areas were quantified using the BZ-X Analyzer software.

### Immunofluorescence staining

Mouse liver sections were deparaffinized and subjected to antigen retrieval, as described above. After blocking with 10% BSA in PBS for 1 h, sections were incubated with primary antibodies overnight at 4°C. After washing with PBS containing 0.1% Tween 20 (PBS-T), sections were incubated with secondary antibodies for 1 h at room temperature, then processed with a Vector TrueVIEW Autofluorescence Quenching Kit (VectorLabs), and nuclei were stained using DAPI. Images were acquired using a BZ-X700 All-in-One fluorescence microscope (Keyence, Osaka, Japan), or an LSM800 confocal microscope (Carl Zeiss, Oberkochen, Germany).

### Single-cell fixed RNA sequencing analysis

Sample preparation: Frozen liver tissues from three mice per group were used for single-cell fixed RNA sequencing. Tissue fixation was performed using a Chromium Next GEM RNA Profiling Sample Fixation Kit (10x Genomics), according to the manufacturer’s instructions. Fixed cells were stored at −80°C in Quenching Buffer supplemented with Enhancer and Glycerol.

Pre-processing: Fixed single-cell suspensions (n = 3 per group) were loaded onto a 10x Chromium system (10x Genomics), and libraries were prepared according to the manufacturer’s instructions. Sequencing was performed on the Illumina NovaSeq 6000 platform. Raw reads were processed using Cell Ranger (v7.1.0, 10x Genomics) against the mm10 mouse reference transcriptome.

Integration and clustering: Data were analyzed using Seurat (v4.3.0.1). Highly variable genes (n = 3,000) were selected using SelectIntegrationFeatures. Datasets were integrated using reciprocal PCA (rpca) *via* FindIntegrationAnchors and IntegrateData. The default assay was set to “integrated,” followed by ScaleData, RunPCA, and ElbowPlot to determine the number of principal components (PCs). Clustering and dimensionality reduction were performed using FindNeighbours, FindClusters, and RunUMAP. Clusters that were visually distinct in UMAP but shared the same identity were manually separated using CellSelector. Reclustering was done using standard procedures (ScaleData, FindVariableFeatures, RunPCA, FindNeighbours, FindClusters, RunUMAP).

Cell type annotation: Differentially expressed genes (DEGs) were identified for each cluster using FindMarkers. Cell types were annotated manually based on known markers or by referencing the top DEGs against the PanglaoDB and CellMarkers databases.

Gene set enrichment analysis (GSEA): Gene ontology (GO) enrichment was conducted using enrichR (v3.2) or ClusterProfiler (v4.6.2), referencing the “GO_Biological_Process_2023,” “GO_Cellular_Component_2023,” and “GO_Molecular_Function_2023” databases. DEGs (log2FC > 0.25, up to 100 genes per cluster) were identified with FindAllMarkers. Representative pathways with significantly adjusted p-values were manually curated.

Cell-Cell Communication Analysis: CellChat (v1.6.1) was used to infer cell-cell communication, based on the CellChatDB ligand-receptor database. Pathway-level networks were constructed by aggregating ligand-receptor interactions and compared across conditions.

### Plasmids

Open reading frames of cDNAs were amplified by reverse transcription-PCR. Mutants of these cDNAs were prepared by the QuikChange method, and the entire coding sequences were verified. The resulting cDNAs were ligated to the appropriate epitope sequences and cloned into the pcDNA3.1 vector (Invitrogen). For lentiviral transduction of *Lrba*, pCSII-CMV-RfA-IRES-Blast (RIKEN BioResource Research Center) was used.

### Cell culture and transfection

HEK293T and mouse embryonic fibroblasts (MEF) were cultured in DMEM, containing 10% fetal bovine serum (FBS), 100 IU/ml penicillin G, and 100 μg/ml streptomycin, at 37°C under a 5% CO_2_ atmosphere. KUP5 (mouse Kupffer cell line)(Kitani *et al*., 2014) cells were cultured in DMEM containing 10% FBS, antibiotics, 10 μg/ml bovine insulin, and 250 μM monothioglycerol.

Bone marrow-derived macrophages (BMDM) were prepared as described(Weischenfeldt & Porse, 2008). Briefly, bone marrow cells were isolated from the femurs and tibias of mice. After depletion of reticulocytes, the residual cells were cultured in RPMI, containing 10% FBS, 5% L-929 conditioned medium, 55 μM 2-mercaptoethanol, and antibiotics, at 37°C under a 5% CO_2_ atmosphere. The floating cells were removed, and the resultant adherent cells were used as macrophages on day 7.

Plasmid transfection into HEK293T cells was performed using polyethylenimine (PEI). Transfection of siRNAs was performed using Lipofectamine RNAiMAX (Thermo Fisher Scientific). For the stable expression of FLAG-tagged *Lrba* in *Lrba*^-/-^ MEFs, lentiviral infection followed by selection with 5 µg/ml blasticidin was performed.

### Construction of *Lrba*^-/-^ cells

To generate *Lrba*^-/-^ MEFs, the gRNA cloning vector (Addgene; #41824) targeting 5ʹ-atgcccctgatggggagcccCGG-3ʹ (PAM sequence in uppercase) in the coding sequence of mouse *Lrba*, and the pCAG-hCas9 expression vector (Addgene; #51142) were transfected using Lipofectamine 3000 (Thermo Fisher). After 96 h, cells were cloned by limiting dilution to obtain single-cell clones. The clones were validated by *Ban*II digestion, sequenced with PCR-amplified fragments to confirm mutations, and subjected to immunoblotting with an anti-LRBA antibody.

### Immunoprecipitation, SDS-PAGE, and immunoblotting

For immunoblotting, liver tissues and cells were lysed with RIPA buffer (50 mM Tris-HCl, pH 7.5, containing 150 mM NaCl, 1% NP-40, 1% sodium deoxycholate, and 0.1% SDS). For immunoprecipitation, cells were lysed with 50 mM Tris-HCl, pH 7.5, containing 150 mM NaCl, 1% Triton X-100, 2 mM PMSF, and a complete protease inhibitor cocktail (Sigma). Immunoprecipitation was performed using appropriate antibodies, followed by Protein G agarose beads (Cytiva) at 4°C with gentle rotation. Immunoprecipitates were washed five times with the lysis solution. Samples were separated by SDS-PAGE and transferred to PVDF membranes. After blocking, the membranes were incubated with the appropriate primary antibodies, followed by an incubation with HRP-conjugated secondary antibodies. Chemiluminescent images were obtained with a Fusion Solo S imaging system (Vilber).

### qPCR

For analyses with mouse tissues, total mRNA was extracted with an RNeasy Mini Kit (QIAGEN) and then transcribed to cDNA with ReverTra Ace qPCR RT Master Mix with gDNA Remover (TOYOBO), according to the manufacturers’ instructions. For analyses with cultured cells, cell lysis and reverse transcription were performed with a SuperPrep Cell Lysis RT Kit for qPCR (TOYOBO). qPCR was performed with Power SYBR Green PCR Master Mix (Life Technologies), according to the manufacturer’s instructions. Quantitative real-time PCR was performed with a CFX Connect Real-Time System (BIO-RAD) by the ΔΔCT method. Primer sequences are listed in Appendix Table S3.

### Identification of LRBA-associated proteins by proximity labeling

HEK293T cells were transfected with plasmids encoding AGIA(Yano *et al*, 2016)- and AirID(Kido *et al*, 2020)-tagged EGFP or LRBA in 10-cm dishes. At 18 h after transfection, the cells were treated with 50 μM biotin for 3 h, washed with HEPES-saline buffer (20 mM HEPES-NaOH, pH 7.5, 137 mM NaCl), and lysed with Gdm-TCEP buffer (6 M guanidine-HCl, 100 mM HEPES-NaOH, pH 7.5, 10 mM TCEP, 40 mM chloroacetamide). After heating and sonication, proteins in the cell lysates (*n = 3* per group) were purified by methanol–chloroform precipitation and solubilized using PTS buffer (12 mM SDC, 12 mM SLS, 100 mM Tris-HCl, pH 8.0). After sonication and 5-fold dilution with 100 mM Tris-HCl, pH 8.0, the protein solutions were digested with trypsin (MS grade, Thermo Fisher Scientific) at 37°C overnight. The digests were diluted 2-fold with TBS (50 mM Tris-HCl, pH 7.5, 150 mM NaCl), and biotinylated peptides were captured on a 15 µL slurry of MagCapture HP Tamavidin 2-REV magnetic beads (FUJIFILM, Wako) by an incubation for 3 h at 4°C. After five washes with TBS, the biotinylated peptides were eluted with 100 µL of 1 mM biotin in TBS for 15 min at 37°C, twice. The combined eluates were desalted using a GL-Tip SDB (GL Sciences), evaporated, and dissolved in 0.1% TFA and 3% acetonitrile (ACN).

LC-MS/MS analysis of the resultant peptides was performed on an EASY-nLC 1200 UHPLC connected to an Orbitrap Fusion mass spectrometer through a nanoelectrospray ion source (Thermo Fisher Scientific). The peptides were separated on a 150-mm C18 reversed-phase column with an inner diameter of 75 µm (Nikkyo Technos), with a linear 4–32% CAN gradient for 0–60 min, followed by an increase to 80% ACN for 10 min. The mass spectrometer was operated in the data-dependent acquisition mode with a maximum duty cycle of 3 s. MS1 spectra were measured with a resolution of 120,000, an automatic gain control (AGC) target of 4 × 10^5^, and a mass range of 375–1500 *m/z*. HCD MS/MS spectra were acquired in the linear ion trap with an AGC target of 1 × 10^4^, an isolation window of 1.6 *m/z*, a maximum injection time of 200 ms, and a normalized collision energy of 30. Dynamic exclusion was set to 10 s. Raw data were directly analyzed against the Swiss-Prot database restricted to *Homo sapiens*, using Proteome Discoverer version 2.4 (Thermo Fisher Scientific) with the Mascot search engine. The search parameters were as follows: (a) trypsin as an enzyme with up to two missed cleavages; (b) precursor mass tolerance of 10 ppm; (c) fragment mass tolerance of 0.6 Da; (d) carbamidomethylation of cysteine as a fixed modification; and (e) acetylation of protein N-terminus, oxidation of methionine, and biotinylation of lysine as variable modifications. Peptides were filtered at a false discovery rate (FDR) of 1%, using the Percolator node. Label-free quantification was performed based on the intensities of precursor ions, using the Precursor Ion Quantifier node. Normalization was performed such that the total sum of abundance values for each sample over all peptides was the same. For statistical analyses of MS data, the P-values in each volcano plot were calculated using Student’s two-sided t-tests.

### Structural prediction and modeling of complexes

Structural prediction and modeling were conducted using the AlphaFold Server powered by AlphaFold 3 (https://alphafoldserver.com/)(Abramson *et al*, 2024). The structures of the full-length LRBA monomer, or the complex of the LRBA PH-BEACH domain with the MAPK family, MAP2K family, MAP3K family, IKKα, IKKβ, IKKγ, IKKε, and TBK1, were predicted, and the interface predicted template modeling (ipTM) scores of the complexes were listed. For those with an ipTM of 0.7 or higher, the structure and Predicted Aligned Error (PAE) were shown.

### Gene expression database analysis

Bulk RNA-sequencing data were retrieved from the Gene Expression Omnibus (GEO) database, under accession number GSE135251(Zeldin, 2001). Expression levels in healthy patients and patients with MASLD expression were analyzed using the GEO2R tool and R statistical software (version 4.2.3).

To investigate the cell-specific expression of LRBA, single-nucleus RNA-sequencing (snRNA-seq) data sets were obtained from GSE185477 (two normal livers), GSE174748 (two normal and two steatotic livers), and GSE212046 (two cirrhotic livers). Raw data were processed with the R (v.4.3.3) statistical software and the Seurat package (v.5.0.3). After conversion into a Seurat object, datasets were integrated using the anchor-based reciprocal principal component analysis (RPCA). Filtering was applied to remove any cells with >15% mitochondrial transcripts, <200 detected genes, or >5,000 total genes. Cell types were annotated based on canonical marker genes. Uniform manifold approximation and projection (UMAP) were used for dimensionality reduction, and *LRBA* expression was visualized using UMAP plots, violin plots, and pie charts.

### Statistics

One-way ANOVA followed by a post-hoc Tukey HSD test, the Mann-Whitney test, and the log-rank test for Kaplan-Meier survival analysis were performed using the GraphPad Prism 10 software. For all tests, a *P* value <0.05 was considered statistically significant.

## Acknowledgements

We thank Ms. Shiori Motoyama, Ms. Chihiro Yamada, Dr. Hiroko Ikejima (Osaka Metropolitan Univ.), and Ms. Yukimi Kira in the Research Support Platform of Osaka Metropolitan University Graduate School of Medicine for technical assistance. This work was partly supported by AMED (JP24gm6410013 to D.O., JP24wm0625516 to F.T.), a Grant for Research Program on Hepatitis from the Japan Agency for Medical Research and Development (19fk0210050 to F.T.), MEXT/JSPS KAKENHI grants (JP24K02242, JP22K18385 to F.T., JP24H01909, JP21H00291 to D.O., and JP22K07174 to K.S.), JST ACT-X Grant Number JPMJAX2117 (K.S.), Suzuken Memorial Foundation (D.O.), Waksman Foundation of Japan (D.O.), Kobayashi Foundation (F.T.), Medical Research Center Initiative for High Depth Omics (H.K.), and Joint Usage and Joint Research Programs of the Institute of Advanced Medical Sciences, Tokushima University (F.T.).

## Author contributions

D.O., Y.O., K.S., and M.S. performed biochemical and cell biological experiments and mouse analyses. M.G. performed histochemical analyses. D.P.M., L.T.T.T., H.I., M.K., and T.M. performed scRNA-seq data analysis. Y.S. performed structural prediction. H.K., H.T., and T.S. conducted proximity labeling and mass spectrometric analyses. F.A. and S.U. provided the LRBA antibody. T.H. and I.H. made the knockout mouse. K.I., N.K., and F.T. conceived the project. D.O. and F.T. wrote the manuscript, and all of the authors participated in commenting on the manuscript.

## Disclosure and competing interests statement

The authors declare no competing interests.

## Data availability

The scRNA-seq data with *Lrba^-/-^* mice have been deposited at the Gene Expression Omnibus (GEO) with accession code GSE301261. The deposited data are publicly available as of the date of publication. In addition, this paper analyzed existing, publicly available data. All data supporting the findings of this study are available within the main text and supplemental material and from the corresponding authors upon request.

---

**Figure EV1.**
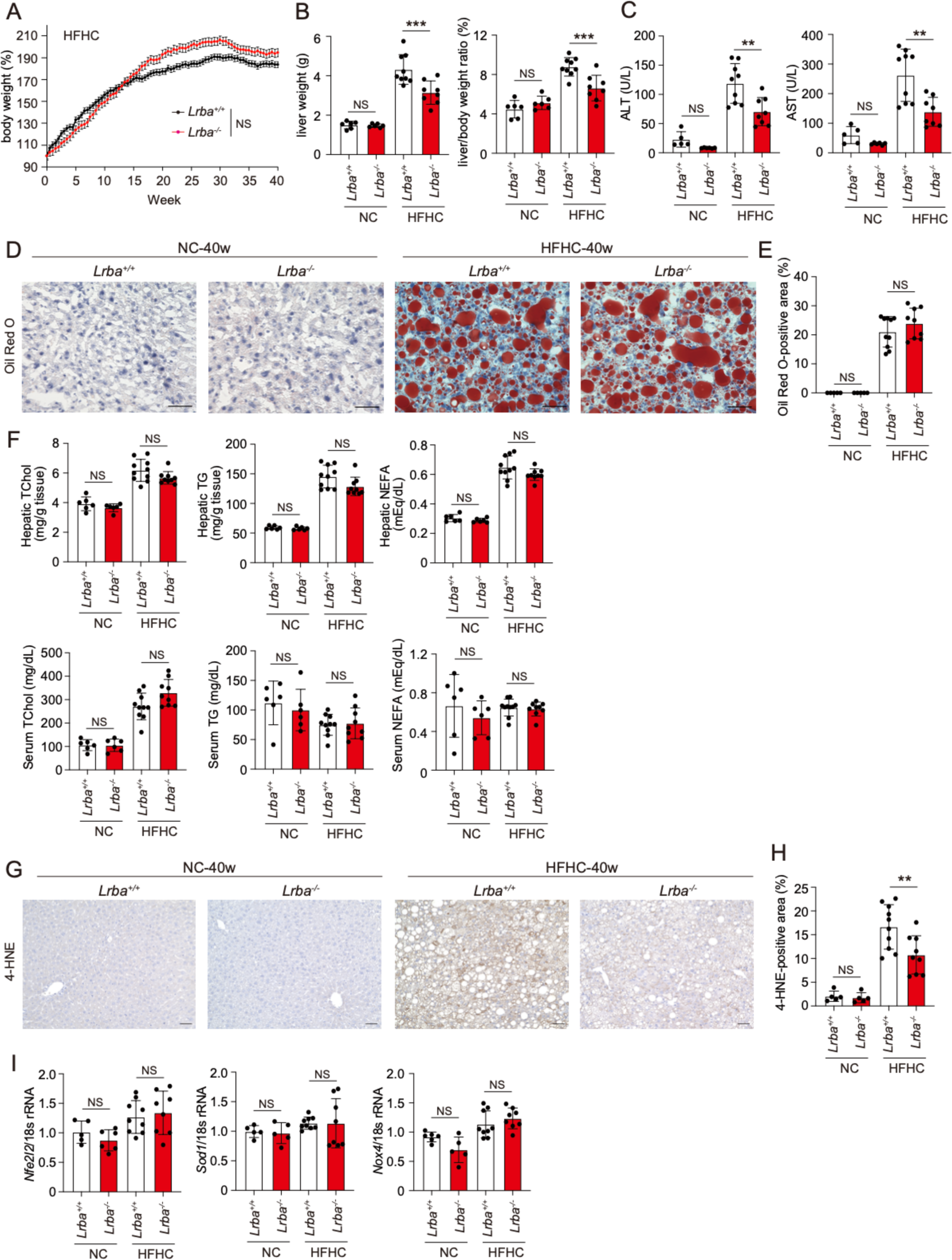
Comparable lipid accumulation in livers from HFHC diet-fed *Lrba*^+/+^ and *Lrba*^-/-^ mice. (**A**) Comparable body weights during HFHC feeding. Body weights of *Lrba*^+/+^ and *Lrba***^-/-^** mice are shown (*n* = 6-10). (**B**) Attenuated hepatomegaly in HFHC-fed *Lrba***^-/-^**mice. Liver weight (left) and liver-to-body weight ratio (right) in NC- or HFHC-fed *Lrba*^+/+^ and *Lrba***^-/-^** mice are indicated (*n* = 6-9). (**C**) Reduced serum ALT and AST levels in HFHC-fed *Lrba***^-/-^** mice (*n* = 5-9). (**D**-**F**) Comparable lipid accumulation between HFHC-fed *Lrba*^+/+^ and *Lrba***^-/-^**mice. Representative liver images stained with Oil Red O (**D**), quantification of Oil Red O-positive area (**E**) (*n* = 5-10), and measurements of total cholesterol (TChol), triglyceride (TG), and non-esterified fatty acid (NEFA) levels in serum or liver tissue samples (**F**) (*n* = 6-10) are shown. (**G**, **H**) Reduced lipid oxidation in HFHC-fed *Lrba***^-/-^** mice. Representative liver sections stained for 4-HNE (**G**) and quantification of 4-HNE-positive area (**H**) (*n* = 5-10). (**I**) Induction of oxidative stress-related genes was not affected in HFHC-fed *Lrba***^-/-^** mice. qPCR analysis of indicated targets was performed using the total liver RNA (*n* = 5-9). Data are shown as mean ±SD by *t*-test (A), or individual values with mean ± SEM. **p < 0.01, ***p < 0.001 by the ANOVA post-hoc Tukey test (B, C, E, F, H, I). NS, not significant.

**Figure EV2.**
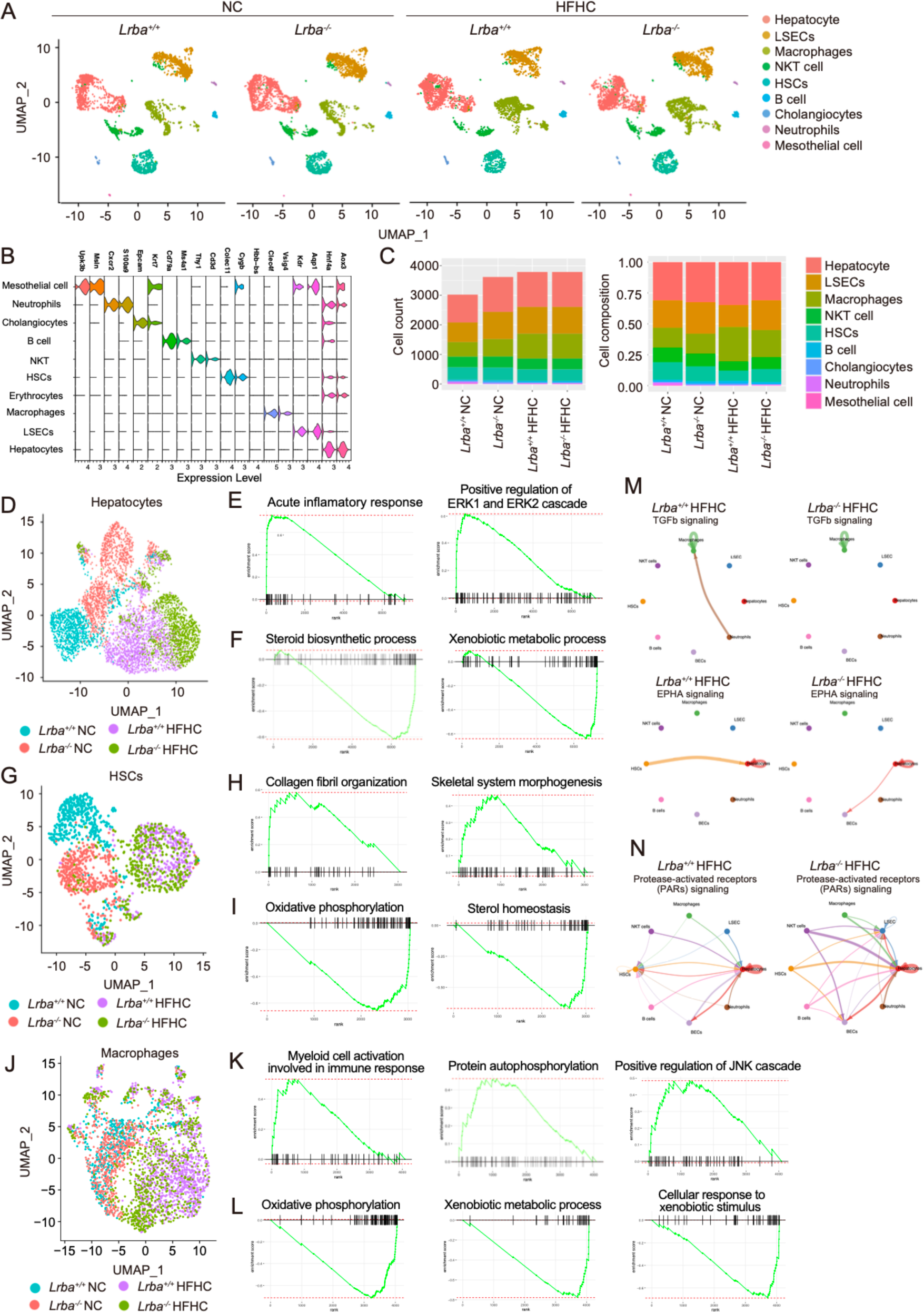
scRNA-seq analysis of the livers from HFHC diet-fed *Lrba*^+/+^ and *Lrba*^-/-^ mice. (**A**-**C**) Whole liver cells were classified into nine distinct clusters. UMAP plots of whole liver cells from NC- or HFHC diet-fed *Lrba*^+/+^ and *Lrba***^-/-^** mice (**A**), cluster-defining marker genes (**B**), and cell count or cell composition (**C**) are shown. (**D**-**F**) Inflammatory responses were attenuated in HFHC-fed *Lrba***^-/-^** mice hepatocytes. A UMAP plot of the hepatocyte cluster (**D**), gene ontology enrichment analysis of pathways downregulated (**E**) or upregulated (**F**) in HFHC-fed *Lrba***^-/-^** mice compared with that of *Lrba*^+/+^ mice are shown. (**G**-**I**), Collagen fibril organization was attenuated in the HSCs from *Lrba***^-/-^** mice. UMAP plot of HSC cluster (**G**), and enrichment analyses of pathways downregulated (**H**) or upregulated (**I**) in HSCs from HFHC-fed *Lrba***^-/-^**mice are shown. (**J**-**L**), Myeloid cell activation, protein autophosphorylation, and positive regulation of JNK cascade were attenuated in the *Lrba*^-/-^ mice macrophages. UMAP plot of macrophage cluster (**J**), and enrichment analyses of downregulated (**K**) or upregulated (**L**) pathways in HFHC-fed *Lrba***^-/-^** mice are indicated. (**M**, **N**) Intercellular communications affected between HFHC-fed *Lrba*^+/+^ and *Lrba*^-/-^ mice. CellChat analysis of intercellular signaling pathways that are attenuated (**M**) or enhanced (**N**) in HFHC-fed *Lrba***^-/-^** mice compared to HFHC-fed *Lrba*^+/+^ mice.

**Figure EV3.**
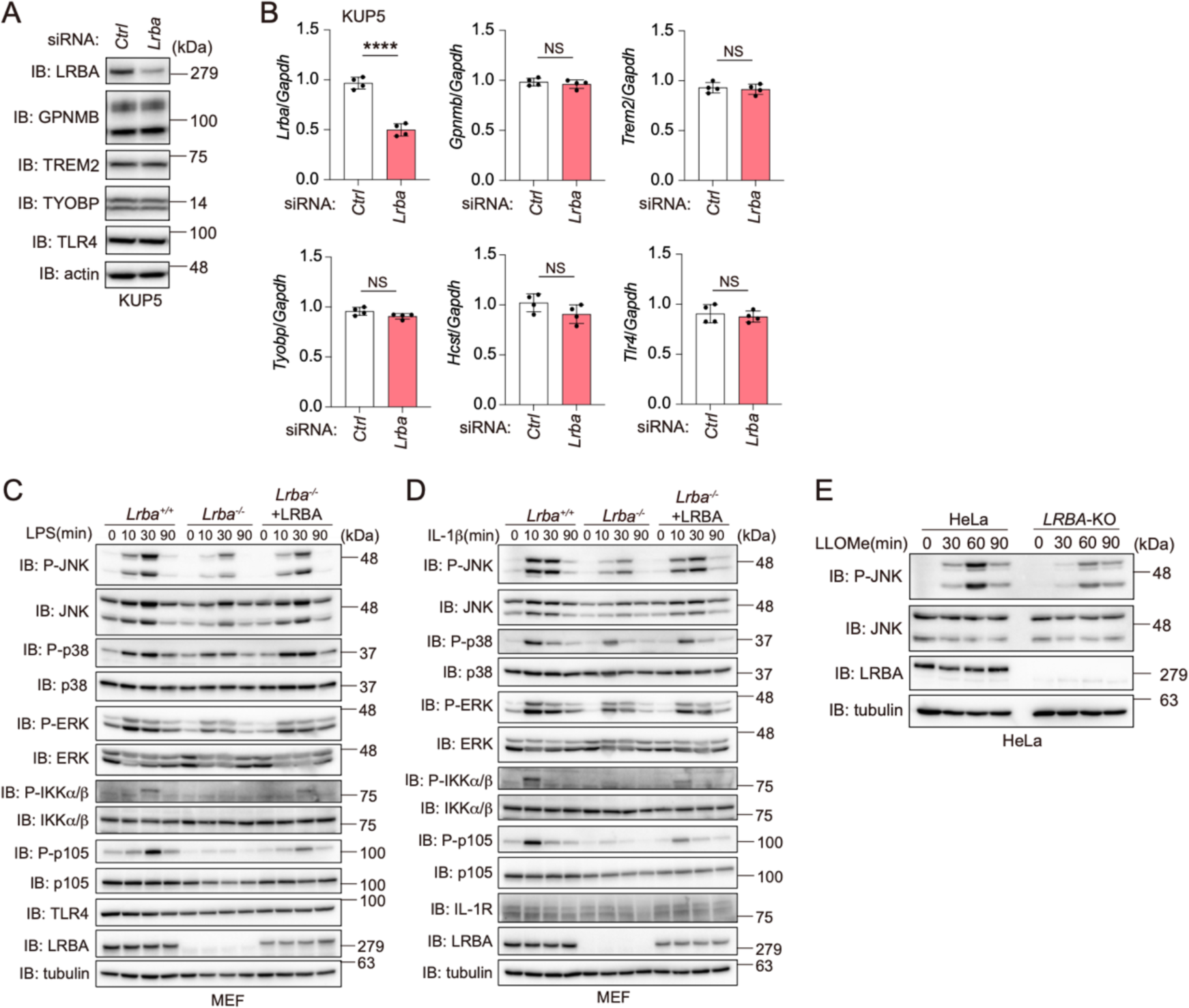
*Lrba*-knockdown attenuates MAPK signaling. (**A**, **B**) KUP5 mouse Kupffer cells were transfected with the indicated siRNAs and analyzed by immunoblotting using the indicated antibodies (**A**) and by qPCR analysis (**B**) (*n* = 4). (**C**-**E**) *Lrba*^+/+^ and *Lrba*^-/-^ MEFs were stimulated with 5 μg/ml LPS (**C**), 20 ng/ml IL-1β (**D**), or 1 mM LLOMe (**E**) for the indicated periods, and the cell lysates were immunoblotted with the indicated antibodies. Data are presented as individual values with mean ± SEM. \*\*\*\**p* < 0.0001 by the Mann-Whitney test (B). NS, not significant.

**Figure EV4.**
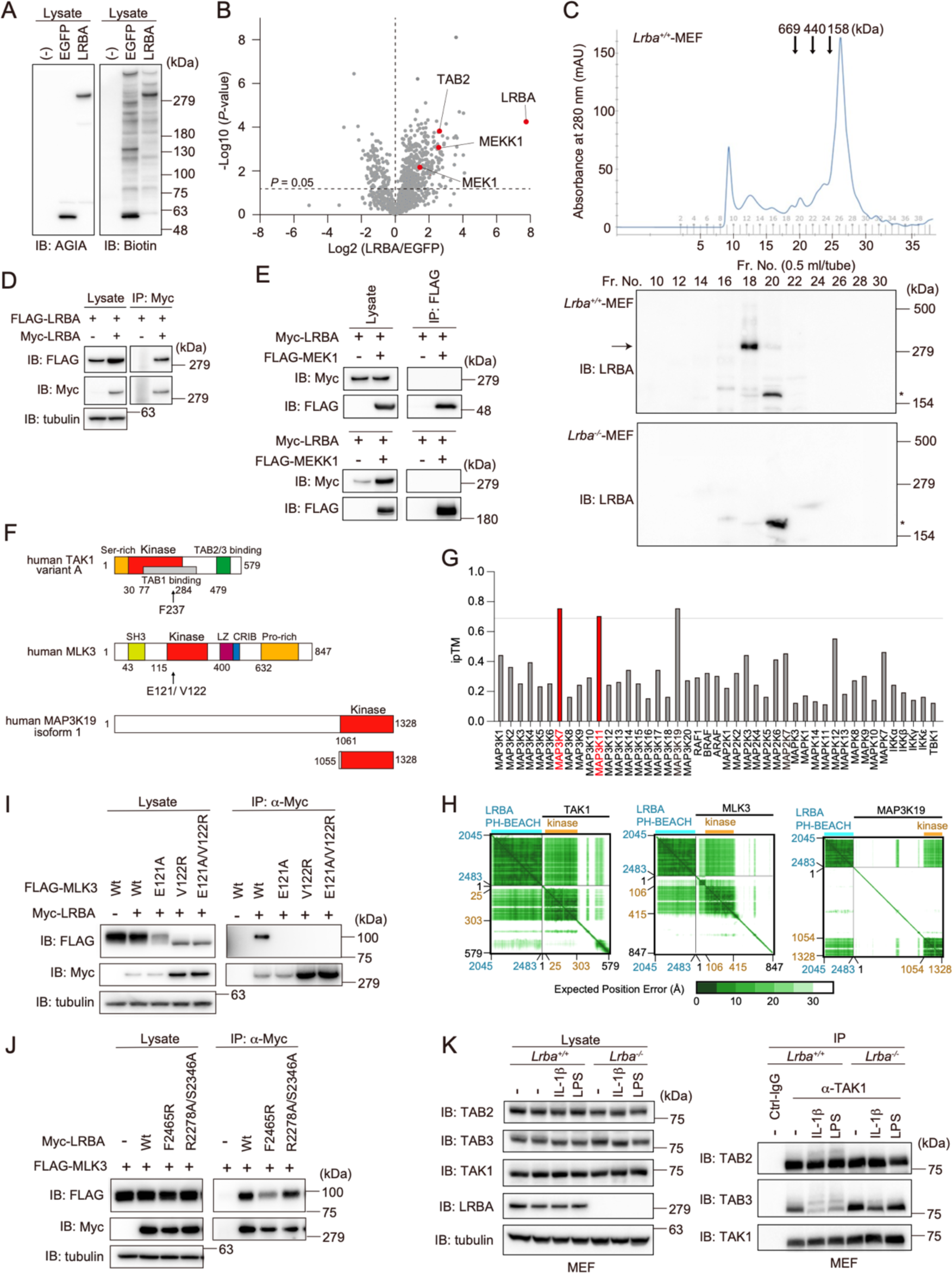
Dimer formation of LRBA and its interaction with MAP3Ks. (**A**) Proximity labeling of LRBA interactors using AirID-LRBA. AGIA- and AirID-tagged LRBA were expressed in HEK293T cells and treated with 50 µM biotin for 3 h. Cell lysates were immunoblotted with the indicated antibodies. (**B**) Identification of candidate LRBA-interacting proteins. Biotinylated peptides were enriched using Tamavidin 2-REV and analyzed by LC-MS/MS. Statistically significant interactors were determined by Student’s two-sided *t*-test in the volcano plots. (**C**) LRBA forms a dimer in cells. Lysates from *Lrba*^+/+^ and *Lrba***^-/-^** MEFs were fractionated using a Superose 6 gel filtration column. Elution profiles from *Lrba*^+/+^ MEFs (*top*), elution of molecular weight standards (*arrows*), and immunoblotting with anti-LRBA antibody (*middle* and *bottom*) are shown. *: nonspecific signal. (**D**) Intermolecular interaction of LRBA. FLAG- and Myc-tagged LRBA were co-expressed in HEK293T cells. Cell lysates and anti-Myc immunoprecipitates were subjected to immunoblotting with the indicated antibodies. (**E**) LRBA does not interact with MEK1 or MEKK1. FLAG-MEK1 or FLAG-MEKK1 was co-expressed with Myc-LRBA in HEK293T cells. Cell lysates and anti-FLAG-immunoprecipitates were immunoblotted. (**F**) Schematic domain structures of TAK1, MLK3, and MAP3K19. Ser-rich: serine-rich domain; SH3: Src homology 3 domain; LZ: leucine-zipper domain; CRIB: Cdc42- and Rac-interactive binding motif; Pro-rich: proline-rich domain. (**G**) AlphaFold 3 screening of MAPK and IKK family kinases for predicted interactors with the LRBA PH-BEACH domain (a.a. 2045-2483). Predicted interaction scores (ipTM) are shown. Experimentally verified interactions are indicated in *red*. (**H**) Predicted aligned error (PAE) plots from AlphaFold 3 for complexes between the LRBA PH-BEACH domain (a.a. 2045-2483) with full-length TAK1, MLK3, or MAP3K19. (**I**) E121 and V122 in MLK3 are required for LRBA binding. A co-immunoprecipitation analysis as in (E) was performed using FLAG-tagged MLK3 Wt or mutants (E121A, V122R, E121A/V122R) with Myc-LRBA. (**J**) Phe2465 in LRBA is important for MLK3 binding. A co-immunoprecipitation analysis as in (E) was performed using Myc-tagged LRBA Wt or mutants (F2465R, R2278A/S2346A) with FLAG-MLK3. (**K**) LRBA does not disrupt TAK1-TAB2/3 interactions. *Lrba*^+/+^ and *Lrba***^-/-^** MEFs were stimulated with 20 ng/ml IL-1β for 10 min or 5 μg/ml LPS for 20 min. Cell lysates and anti-TAK1 immunoprecipitates were subjected to immunoblotting with the indicated antibodies.

**Figure EV5.**
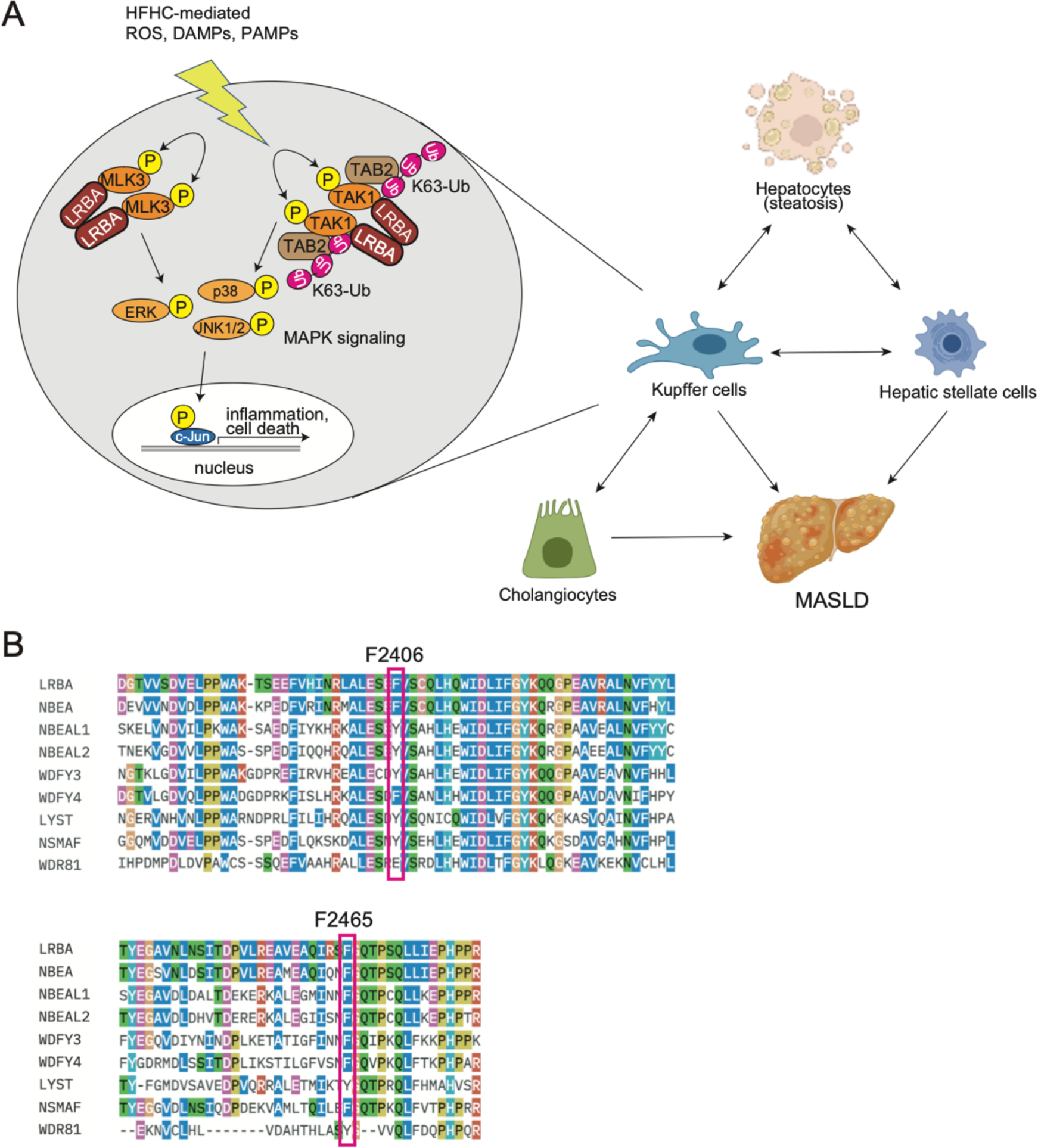
Schematic model for the involvement of LRBA in MASLD. (**A**) Schematic representation of the participation of LRBA in MASLD. (**B**) Alignment of amino acid sequences of BDCPs and conservation of residues corresponding to F2406 and F2465 in LRBA.

